# Unmasking Crucial Residues in Adipose Triglyceride Lipase (ATGL) for Co-Activation with Comparative Gene Identification-58 (CGI-58)

**DOI:** 10.1101/2023.06.26.546551

**Authors:** Natalia Kulminskaya, Carlos Francisco Rodriguez Gamez, Peter Hofer, Ines Kathrin Cerk, Noopur Dubey, Roland Viertlmayr, Theo Sagmeister, Tea Pavkov-Keller, Rudolf Zechner, Monika Oberer

**Affiliations:** Institute of Molecular Biosciences, University of Graz, Graz, Austria; BioTechMed Graz, Austria; BioHealth Field of Excellence, University of Graz, Graz, Austria

**Keywords:** ATGL, adipose triglyceride lipase, PNPLA2, CGI-58, comparative gene identification-58, ABHD5, co-activation, triacylglycerol hydrolase activity, protein-protein interaction, lipolysis, AlphaFold, protein structure

## Abstract

Lipolysis is an essential metabolic process that releases unesterified fatty acids from neutral lipid stores to maintain energy homeostasis in living organisms. Adipose triglyceride lipase (ATGL) plays a key role in intracellular lipolysis and can be co-activated upon interaction with the protein comparative gene identification-58 (CGI-58). The underlying molecular mechanism of ATGL stimulation by CGI-58 is incompletely understood. Based on analysis of evolutionary conservation, we used site directed mutagenesis to study a C-terminally truncated variant and full-length mouse ATGL providing insights in the protein co-activation on a per-residue level. We identified the region from residues N209-N215 in mouse ATGL as essential for co-activation by mouse CGI-58. ATGL variants with amino-acids exchanges in this region were still able to hydrolyze triacylglycerol at the basal level and to interact with CGI-58, yet could not be activated by CGI-58. Our studies also demonstrate that full-length mouse ATGL showed higher tolerance to specific single amino acid exchanges in the N209-N215 region upon CGI-58 co-activation compared to C-terminally truncated ATGL variants. The region is either directly involved in protein-protein interaction or essential for conformational changes required in the co-activation process. Three-dimensional models of the ATGL/CGI-58 complex with the artificial intelligence software AlphaFold demonstrated that a large surface area is involved in the protein-protein interaction. Mapping important amino acids for co-activation of both proteins, ATGL and CGI-58, onto the 3D model of the complex locates these essential amino acids at the predicted ATGL/CGI-58 interface thus strongly corroborating the significance of these residues in CGI-58 mediated co-activation of ATGL.

## Introduction

Intracellular lipolysis, the breakdown of stored triacylglycerol (TG) is essential to maintain energy homeostasis. Adipose triglyceride lipase (ATGL), also termed PNPLA2, iPLA2 and desnutrin, plays an important role in lipolysis by hydrolyzing long chain FA-containing TGs [1–3]. In addition to the hydrolytic function, ATGL also exerts a transacylation function, which generates TG or fatty acid (FA) esters of hydroxy fatty acids (FAHFAS) by esterification of TG-derived FAs with hydroxy-fatty acids [3–6]. The hydrolytic activity is stimulated at the protein level by the co-activator protein CGI-58 (comparative gene identification-58), also named ABHD5 (Abhydrolase domain-containing protein 5) [5,7]. The stimulatory effect of CGI-58 on ATGL is observed upon direct interaction of both proteins and demonstrates the highest efficiency at approximately equimolar concentrations of enzyme and activator protein [7]. ATGL is inhibited by the proteins G0/G1 switch gene 2 (G0S2), hypoxia inducible lipid droplet associated protein (HILPDA) and the very recently described microsomal triglyceride transfer protein (MTP) [8–17]. It is assumed that CGI-58 and G0S2 interact at different binding sites: Lack of ATGL-activity in G0S2 expressing cells could not be rescued with increased CGI-58 concentrations [18]. Furthermore, ATGL/G0S2 binding was not affected by CGI-58 in co-immunoprecipitation experiments [18]. G0S2 and long-chain acyl-coenzyme A inhibit ATGL’s TG-hydrolytic activity in a non-competitive manner, and the inhibitory effect of oleoyl CoA an ATGL’s TG hydrolytic activity was not alleviated by CGI-58 [19,20]. Together, these data suggest that different mechanism and interfaces govern the activity and regulation of the enzyme.

The physiological relevance of ATGL was thoroughly studied in different genetic knockout and transgenic mouse models: Mice lacking ATGL exhibit increased TG deposition in many tissues and disrupted signaling pathways leading to disturbed energy homeostasis [21,22]. A comprehensive record of published ATGL mouse models and associated phenotypes was rigorously summarized elsewhere [23]. Human patients with mutations in the gene coding for ATGL suffer from TG accumulation in leukocytes and multiple tissues in addition to severe cardiomyopathy [22]. Interestingly, mutations in the human gene of the ATGL co-activator CGI-58 cause severe hepatic steatosis and systemic TG accumulation that is always associated with ichthyosis [7,24]. Analogously, global CGI-58 knockout mice suffer from a lethal skin permeability barrier defect due to impaired ω-O-acylceramide synthesis [21,25–27].

Atomic resolution structures of both proteins, CGI-58 and ATGL, are not available yet. Mouse ATGL (mATGL, 486 amino acids) harbors a “patatin-like phospholipase domain (PNPLA)” within resides I10-K179 (InterPro IPR002641), which is name-giving for all PNPLA-family members [28]. The TG-hydrolytic activity of ATGL is catalyzed by a catalytic dyad, formed by S47 and D166, and residues G14-G19 forming the oxyanion hole [29]. Residues I10-G24 are suggested to be involved in TG binding [30]. *In vitro* studies showed increased lipolytic activity of the truncated variants of ATGL (truncated after D288 or L254 in mATGL) compared to the wild-type enzyme [5,10]. Nevertheless, the exact roles of the C-terminal half of ATGL (residues P260-C486) with respect to lipid droplet (LD) localization and autoregulatory function remain to be established.

Mouse CGI-58 (mCGI-58, 351 amino acids) is a member of the α/β-hydrolase-fold containing protein family comprising of an N-terminal region and an α/β-hydrolase core domain with a cap [31]. In contrast to other α/β-hydrolase protein family members, CGI-58 does not exhibit hydrolytic activity due to the lack of a catalytic nucleophile. Previously, we have shown that the N-terminal, Trp-rich region of CGI-58 is important for LD anchoring and largely disordered [32–34]. Removal of 30 N-terminal amino acids of mCGI-58 disrupted both its ability to localize to LDs and its ability to co-activate ATGL. However, this LD anchor by itself lacks the ability to activate ATGL, indicating that other regions of CGI-58 are necessary for ATGL co-activation [33]. Subsequent mutagenesis studies of CGI-58 (ABHD5) demonstrated a crucial function for residues R299, G328 and D334 of CGI-58 in ATGL co-activation [16,35].

In the current study, we identified evolutionary less conserved parts of ATGL by comparing mammalian ATGL with different phyla in the Kingdom Animalia. Importantly, ATGL variants at positions N209, I212 and N215 exhibited intact or marginally reduced basal activity, however drastically reduced activatability by CGI-58 providing evidence that these residues play a significant role in the co-activation process. Artificial intelligence-based modelling approaches for the three-dimensional (3D) structure of the ATGL/CGI-58 complex also predicted the region comprising N209, I212 and N215 to be involved in protein-protein interaction between enzyme and co-activator. Residues R299, G328 and D334 of CGI-58, are also located in the predicted ATGL/CGI-58 binding interface. In the absence of experimental complex structures, our data provide a good working model of the ATGL/CGI-58 complex.

## Results

### Evolutionary conservation provides an initial rationale for generating ATGL variants

Regulation of ATGL activity on a protein and activity level is very well studied with respect to the interaction of ATGL with CGI-58, G0S2 and HILPDA [10–16,21,22]. Interestingly, the regulatory proteins are not conserved in all species, e.g., the inhibitory protein G0S2 only exists in vertebrates while BLAST-searches of ATGL also reveal proteins with significant hits in non-vertebrates [15,36]. Therefore, we performed an unbiased computational screen to identify amino acids within the sequence of ATGL that mediate its interaction with regulatory proteins. We were primarily interested in the N-terminal half of ATGL since previous *in vitro* studies demonstrated that the truncated ATGL variants M1-L254 or M1-D288 can be activated by CGI-58 or inhibited by G0S2 and HILPDA [7,9,15,19].

We compared the amino acid sequences of ATGL from different species, within the subgroup of ‘mammalia’ and within a larger general group termed ‘animalia’ including different chordates (fish, reptiles, birds, mammals) insects, nematodes and mollusks. The analysis was carried out using the ConSurf software, a designated tool to identify functional regions in proteins by exploiting evolutionary data (Figure 1A) [37]. We hypothesized that catalytically and structurally essential regions are highly conserved, whereas regulatory regions might be i) less conserved when comparing mammalian ATGL with ATGL of non-mammalian species and ii) surface exposed to enable protein-protein interaction.

As shown in Figure 1, the levels of conservation are quite different. Both groups, ‘mammalia’ and ‘animalia’ show large sequence variations in regions P3-K8, V57-C61, G96-T101, T158-Q160 and K179-N180. In line with our general hypothesis, the most conserved region even within ‘animals’ is in the central core and catalytic region of the protein, whereas surface exposed parts of α-helices show higher sequence variability (Figure 1B). Based on our hypothesis (i) on sequence conservation, we identified regions F35-A40, V71-N89, V150-V165, F187-D197, H203-K229 and Y242 to match our criteria of conservation within ‘mammalia‘, yet high diversity within ‘animalia‘. To further narrow down residues of interest, we tested these residues for criteria (ii) by plotting them on a 3D model of ATGL and analyzed the residues with respect to surface exposed residues. Accordingly, we decided to introduce the following residues for introduction of single-amino acid exchanges: L81A, L84A, Y151A, Y164A, F187A, S188A, I193A, L205A, N209A, I212A, I212S, N215A, L216A, Y220A, R221A, L226A, F227A, Y242A. These residues are predominately surface-exposed and cluster on an almost continuous surface area of the protein (Figure 1C). The amino acid exchanges were introduces in the C-terminally truncated variant of mouse ATGL (mATGL288) [5,6,15,20,36]. We predominantly mutated aromatic and aliphatic amino acids residues since these side-chains are frequently involved in biological interactions, whereas small alanine residues typically contribute very little to protein-interactions. The exchange I212S alters the physicochemical properties more drastically by introducing a small, polar side-chain.

**Figure 1:**
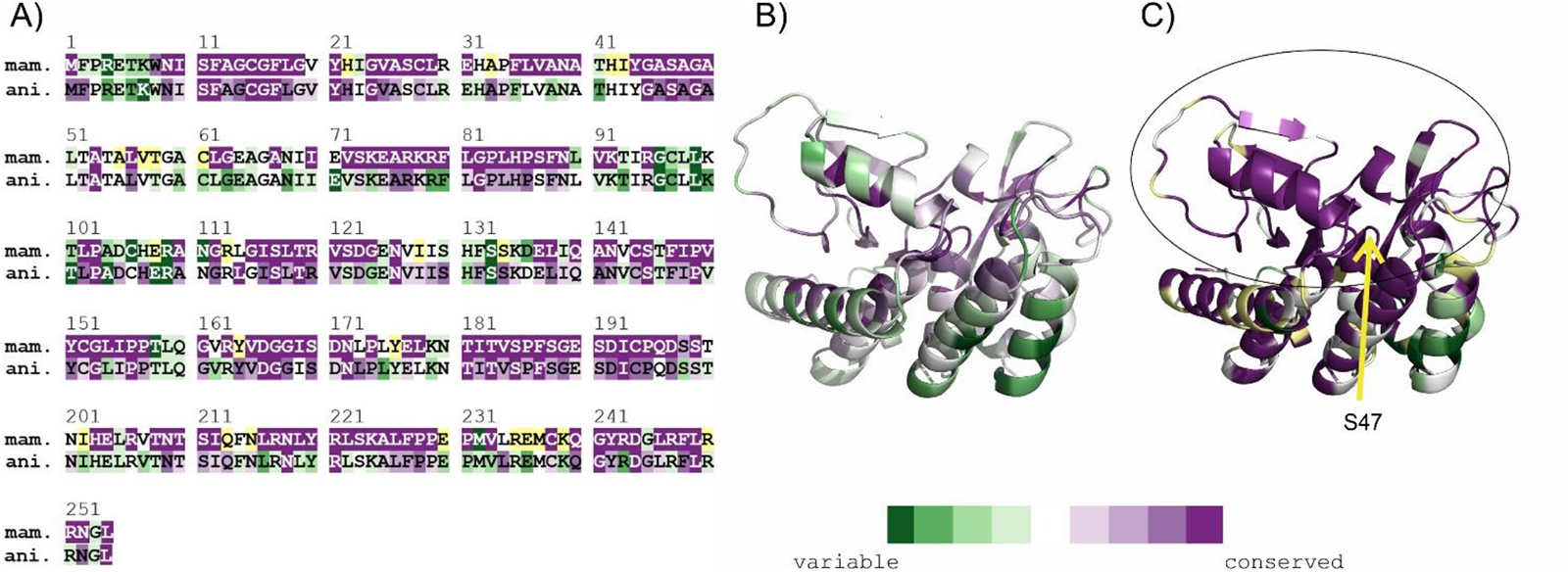
Evolutionary conservation of the minimal variant of mouse ATGL (mATGL254). The rate of conservation is color-coded dark-green (variable) to violet (conserved). Residues with uncertainty in the conservation score are color-coded yellow. A) ConSurf analysis on a per-residue basis within the subgroup of mammals (mam.) compared to the conservation within animals (ani.) in general. B-C). ConSurf analysis plotted on the 3D model of mATGL254 represented in cartoon representation for (B) animals and (C) mammals. The position of the catalytic S47 is indicated with an arrow in (C). The prominent surface cluster of evolutionarily conserved sites in mammals is highlighted with an oval.

### While some ATGL variants retain TG-hydrolytic activity and can be fully co-activated by CGI-58 others lose enzymatic activity and activatability

Next, we expressed each ATGL288 variant in bacterial ArcticExpress (DE3) cells and tested the resulting lysates for TG hydrolase activities in absence or presence of recombinant CGI-58 (Figure 2A, 2C, 2F). The expression levels of ATGL variants in transfected bacterial cells were assessed by immunoblot analyses (Figure 2B, 2D, 2F). To clearly show the difference between basal activity and activatability, we give the X-fold multiplier by which activity is increased upon addition of CGI-58. The multiplier is also insensitive to slight changes in the expression level of ATGL, since basal activity and activity upon CGI-58 stimulation are equally affected. Amino acids F17 and at the active site serine S47 are key amino acids for ATGL activity and the corresponding exchanges F17A and S47A served as negative controls [7,38,39].

We found that ATGL variants carrying the single amino acid substitutions F17A, S47A, Y164A, F187A and I193A exhibit neither basal nor stimulated enzymatic activity (“inactive ATGL variants”) (Figure 2A). Our results also showed that ATGL variants carrying the single amino acid substitutions L81A, L84A, S188A, L205A, L216A, Y220A, R221A and F227A had similar basal and stimulated TG hydrolase activities as wild-type (WT) ATGL (“active ATGL variants”) (Figure 2C). The variants Y151A, L226A and Y242A exhibited significantly reduced but measurable basal activity compared to WT mATGL288 (“partially active ATGL variants”) (Figure 2E).

**Figure 2.**
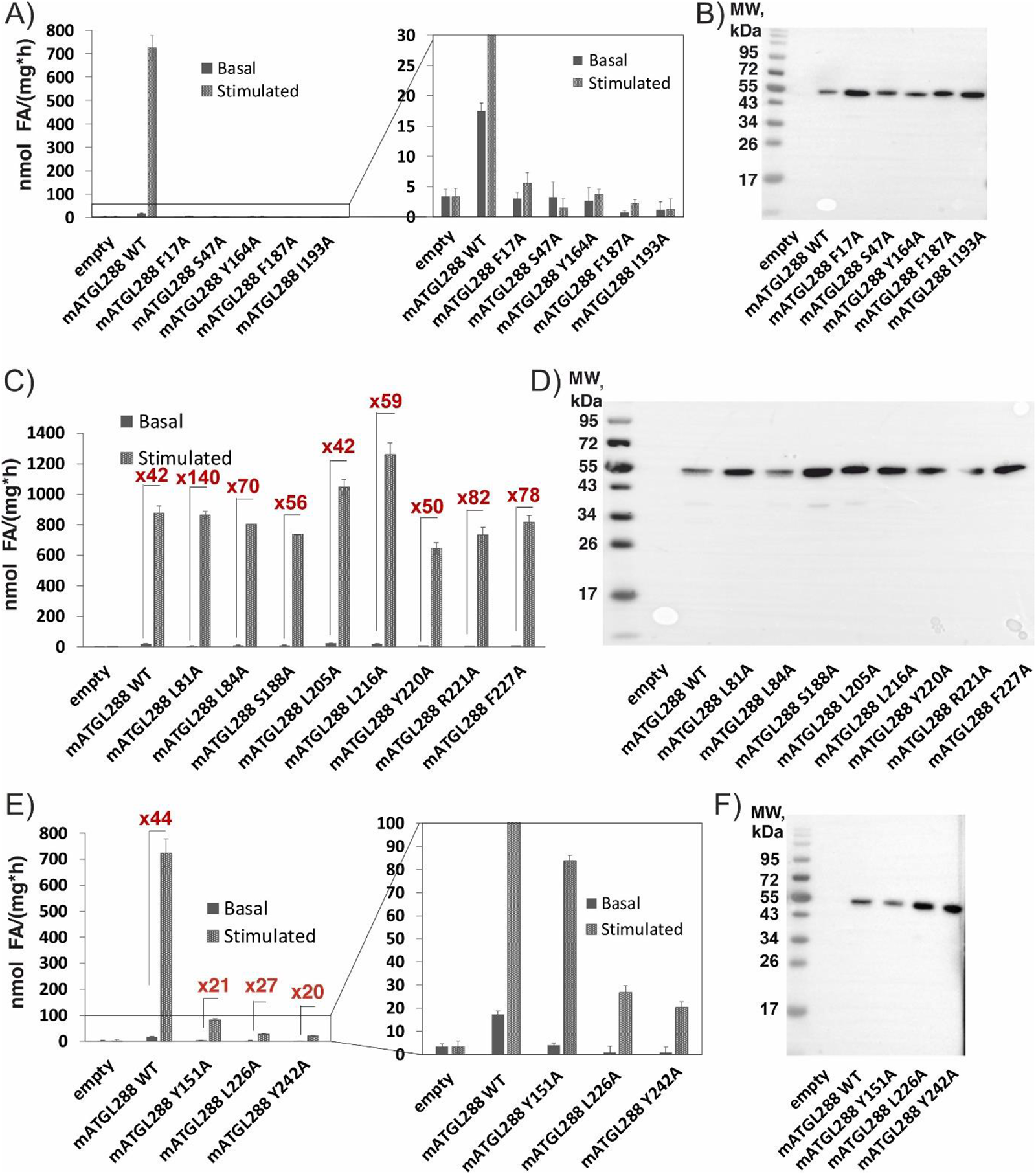
TG hydrolase activity assays and expression control of ATGL variants from bacterial lysates. mATGL288 WT enzymatic activity in basal and mCGI-58 co-activated condition was used as positive control; the multiplier for x-fold increase in activity is indicated for each variant. A) Inactive variants of mATGL288, namely F17A, S47A, Y164A, F187A and I193A in comparison to WT mATGL288 under basal and mCGI-58 co-activated conditions based on enzymatic activity with zoom-in (right). B) Immunoblotting of the expression levels of inactive variants was used as expression control. C) Active variants of mATGL288 namely L81A, L84A, S188A, L205A, L216A, Y220A, R221A and F227A under basal and mCGI-58 co-activated conditions. D) Immunoblotting analysis of the active variants. E) Partially active variants with residual enzymatic activity of mATGL288 namely Y151A, L226A and Y242A; F) Immunoblotting of the expression levels of the partially active variants in comparison to WT mATGL288. Each TG hydrolase activity assay represents the assay of three technical replicates. At least 2 biological replicates were performed.

### ATGL variants N209A, I212A, I212S and N215A have intact basal activity, but lack effective co-activation by CGI-58

When we tested the N209A, I212A, I212S and N215A variants, we observed intact or only slightly reduced basal activity, but reduced activability by mCGI-58 (approximately 1.5 – 4-fold) when compared to activation of WT mATGL288 by mCGI-58 (Figure 3A). To determine the half-maximal effective concentration (EC50) of mCGI-58 to activate ATGL, we performed dose-dependent activity measurements of mATGL288 WT, N209A, I212A, I212S and N215A with increasing concentrations of mCGI-58. The results revealed EC50 values in the range of 500 ± 50 nM for the N209A and I212A variants and 200 ± 20 nM for the I212S and N215A variants, in comparison to EC_50_ values of 178 ± 20 nM for WT mATGL288 (Figure 3B). Immunoblots of the employed bacterial lysates indicated similar expression levels for WT ATGL and the ATGL variants (Figure 3C).

**Figure 3.**
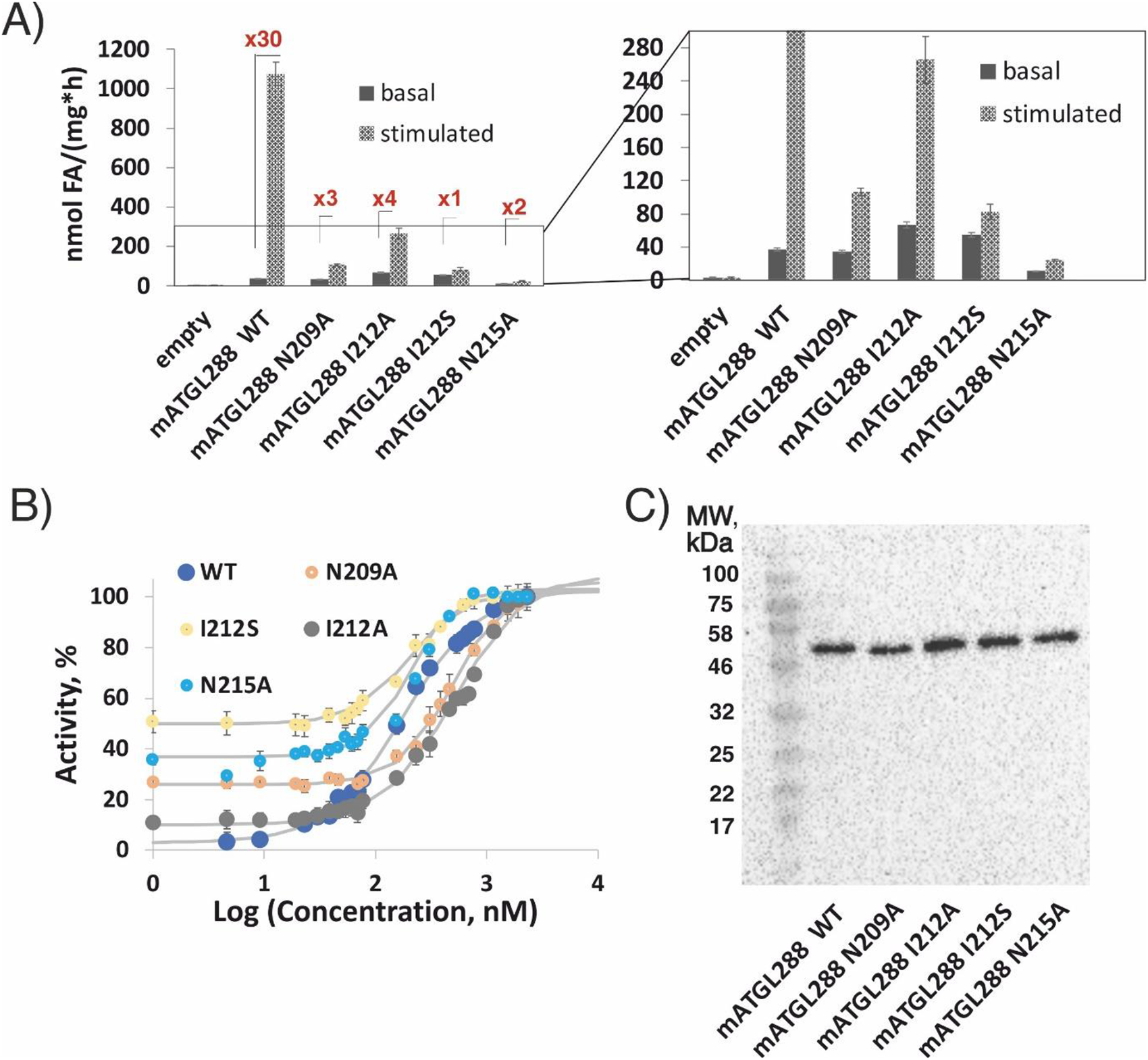
TG hydrolase activity assays of mATGL288 variants that cannot be effectively stimulated by mCGI-58. A) TG hydrolase assays of mATGL288 WT, N209A, I212A, I212S and N215A under basal and mCGI-58 stimulated conditions from bacterial lysates. B) Dose dependent co-activation of TG hydrolytic activity of mATGL288 variants by mCGI-58. Purified mCGI-58 protein was mixed with 25 μg mATGL lysate and tested in TG hydrolytic assays with different mCGI-58 concentrations. 100% activity upon co-activation with 2300 nM CGI-58 corresponds to 925 ± 30 nmol FA/(h*mg) for mATGL288 WT, to 125 ± 4 nmol FA/(h*mg) for mATGL288 N209A; to 352 ± 28 nmol FA/(h*mg) for mATGL288 I212A, to 93 ± 10 nmol FA/(h*mg) for mATGL288 I212S and to 60 ± 5 nmol FA/(h*mg) for mATGL288 N215A. Data are presented as mean ± SD and are representative for three independent experiments. C) Expression controls of mATGL288 WT, N209A, I212A, I212S and N215A were performed using immunoblot analysis. All ATGL variants were tested as bacterial lysates.

### Plotting single amino acid exchanges on a 3D model of mATGL: N209, I212 and N215 are located on a surface region on one face of the protein

To locate the most crucial amino acid residues within the structure of ATGL, we generated a 3D model of mouse ATGL254 using AlphaFold (Figure 4). It is important to note that the model confidence varies significantly between different regions of the enzyme [5,40]. Almost the entire PNPLA domain and two short additional strands of the central β-sheet are modeled with very high confidence, namely residues W8–S73, I94–L173 and T181–P195 (confidence score ‘high’, >90). Similarly, P231–N252 form a long α-helix on the surface of mATGL254 that is predicted with very high confidence. In contrast, residues Q196–E230, which also show largest sequence variability (see Figure 1A), are only predicted with ‘low’ to ‘medium’ confidence scores between 50 and 90. The residues N209, I212 and N215 are located within a region forming a short β-sheet on the surface of ATGL; amino acid exchanges of those residues resulted in retained basal activity yet substantial loss of activability by CGI-58 (Figures 3, 4). Based on the 3D model of ATGL, this region forms a part of the substrate binding pocket that is distant from the catalytic site. We speculate that this region is not directly involved in the catalytic reaction at the scissile ester bond of the lipid, but plays a role in binding or release of the substrate or product (Figure 4).

Substitution of the conserved aromatic residues F17, F187 and Y164 to alanine residues resulted in loss of ATGL-activity which could not be rescued by addition of CGI-58. Amino acid exchanges Y151A, L226A, Y242A resulted in ATGL-variants with reduced basal activity and reduced co-activation upon addition of CGI-58. In the 3D model of mATGL254 residues F17, Y151, F187, L226 are located in or at the entrance to the substrate binding pocket and might therefore not tolerate substitutions (Figure 4). Y164 is positioned in a loop that helps in positioning of the catalytic residue D166 spatially close to S47. Thus, it might be essential for the formation of a principally functioning catalytic active site architecture within ATGL. Y242 is located on the surface exposed face of an α-helix. The complete loss of activity and co-activation upon introducing the change I193A might result from destabilization of ATGL due to loss of hydrophobic interactions (Figure 2 and Figure 4).

**Figure 4:**
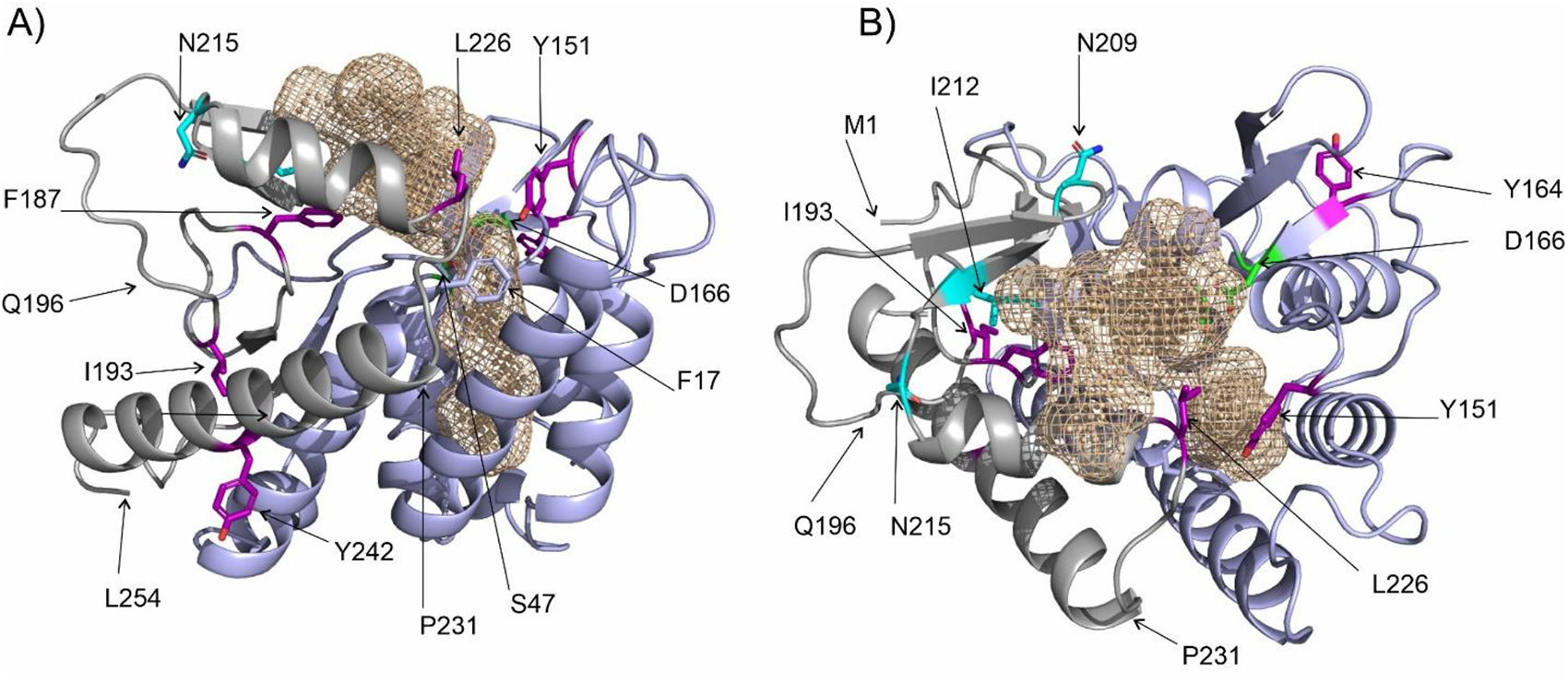
3D model of ATGL depicting important residues for activity and co-activation. The 3D structure of mouse ATGL (M1-L254) is in cartoon representation with the substrate binding cavity as a beige-colored mesh. The annotated PNPLA-domain (I10-K179), which is predominantly predicted with very high confidence, is colored light blue; residues M1-N9 and K179-L254 are in gray. The region (Q196-P231) is involved in shaping the entrance to the active site, yet is predicted with less confidence than the PNPLA-domain. Active site residues (S47, D166) are depicted as green sticks. A) Side-chains of Y151, Y164, I193, L226, Y242 are shown as purple sticks; N209, I212 and N215 as cyan sticks; F17 as light blue stick. The position of L254 is indicated. B) Cartoon representation of mATGL254 as in (A) after approx. 90° rotation to highlight the position of residues N209, I212 and N215 colored as cyan sticks. The positions of M1, Q196 and P231 are indicated with arrows.

### Experiments with full-length ATGL confirm the relevance of N209A, I212A and N215A for enzyme activation by CGI-58

To investigate whether our results with bacterially expressed mATGL288 can be recapitulated with mATGL variants in lysates from a eukaryotic expression system, we transfected the suspension-adapted human embryonic kidney cell line Expi293F with WT and mutated mATGL variants and tested cell lysates for TG-hydrolytic activity (Figure 5). Upon mCGI-58 addition, truncated WT mATGL288 was activated 11-fold, whereas the mATGL288 variants harboring double or triple amino acid exchanges, were only stimulated up to 2-fold (Figure 5A). All mATGL288 variants showed similar expression efficacy in transfected Expi293F cells (Figure 5B). In the eukaryotic system, we also tested full-length mATGL to confirm the results obtained with the truncated mATGL288 enzyme. Addition of CGI-58 increased TG hydrolase activity of lysates containing WT full-length ATGL approximately 20-fold. The single amino acid exchange variants N209A, N212A and N215A were also activatable although to a lesser degree (8 and 14-fold). In contrast, double (N209A/N215A) or triple mutations (N209A/I212A/N215A) in full-length mATGL essentially lost the ability to be co-activated activated by mCGI-58 (Figure 5A)) despite similar expression level (Figure 5B).

**Figure 5.**
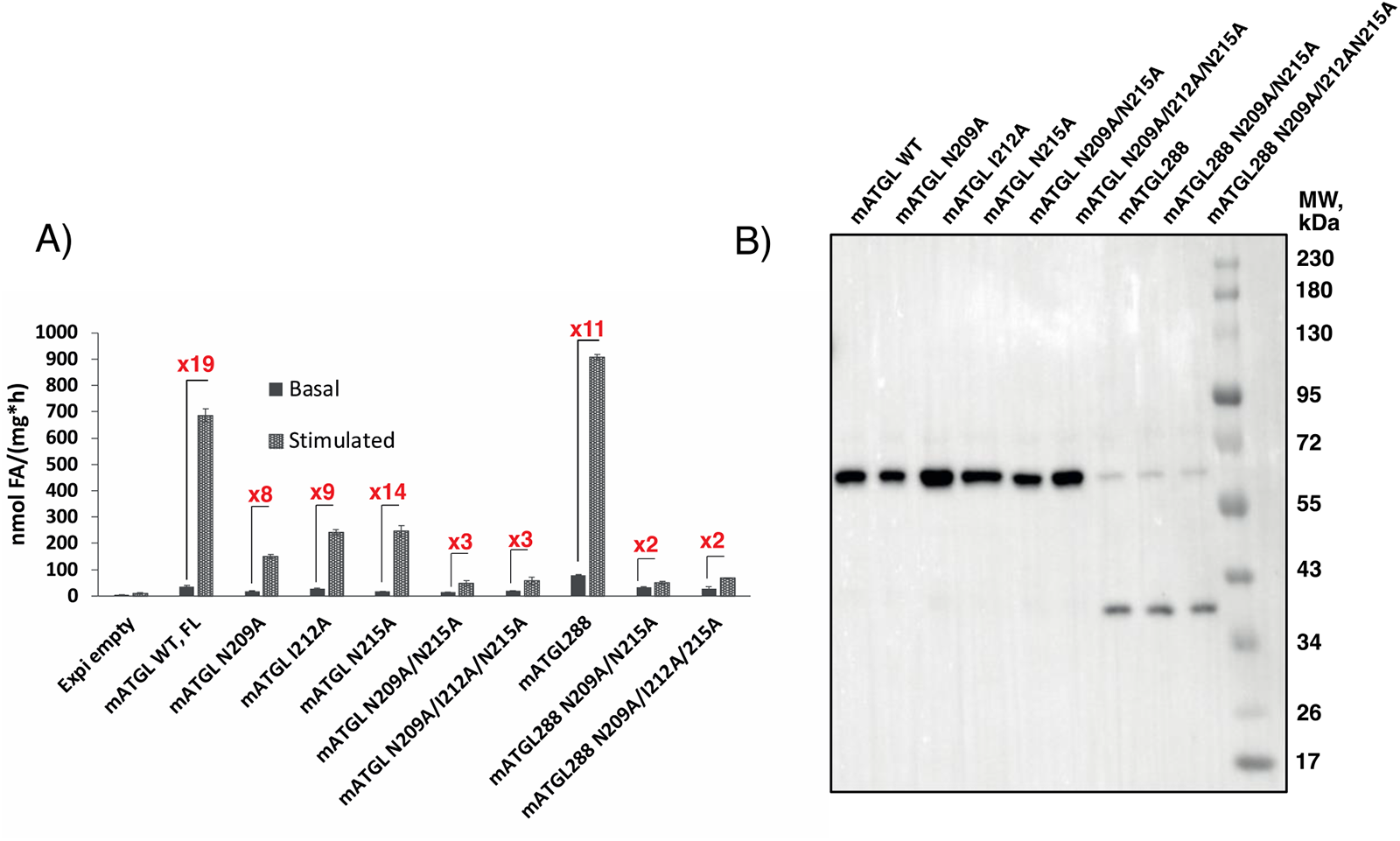
TG hydrolase activity assay of the ATGL variants expressed in mammalian Expi293F cells. A) Basal and mCGI-58 co-activated hydrolytic activity of mATGL in full-length background and in truncations after residue 288 (mATGL288) with single, double and triple amino acid exchanges N209A, I212A, N215A, N209A/N215A and N209A/I212A/N215A. B) Protein expression control of the mATGL variants was performed using immunoblot analysis.

### ATGL variants that cannot be co-activated by CGI-58 preserve the ability to bind CGI-58

Next, we addressed whether loss of activability of the mATGL288 variants N209A, I212A, I212S, N215A is a consequence of the enzyme’s inability to interact with mCGI-58. Therefore, we used the pST44 polycistronic expression system for co-expression of StrepII-tagged mATGL288 (smt3-mATGL288-StrepII) and variants thereof (N209A, I212A, I212S, N215A and N209A/I212A/N215A) together with His-tagged mCGI-58 (smt3-mCGI-58-His6) (Figure 6A). An analogous pST44 polycistronic expression system has been used before for testing ATGL/G0S2 interaction [36].

After cell lysis, we employed affinity purification, to isolate mATGL288 and potential mATGL288/mCGI-58 protein-protein complexes. Lysates and column fractions were analyzed for complexed and uncomplexed mATGL288 and CGI-58 by Western blotting analyses using an anti-StrepII-antibody and anti-His-antibody for the detection of mATGL variants or mCGI-58, respectively (Figure 6B) : cell lysates (Lys) were analyzed to demonstrate co-expression of the proteins; flow through (FT) to check for unbound protein; the last wash fraction (W) to verify that all unbound proteins had been removed after loading to the StrepII affinity column. Finally, two elution fractions (EI and EII) were tested for the presence of mATGL288/mCGI-58 complexes. Despite relatively low mATGL288-StrepII-expression levels in the lysates, mATGL288-StrepII variants efficiently bound to the StrepII column and were easily detected upon elution (Figure 3B, left). While mATGL288-StrepII was retained by the Strep column, uncomplexed His-tagged mCGI-58 expectedly passed through the StrepII-column (Figure 3B, right). After extensive washing to remove all uncomplexed mCGI-58-His, StrepII-tagged mATGL288 and its complexes were eluted from the StrepII column with desthiobiotin . When we co-expressed the mATGL288 variants N209A, I212A, I212S, N215A and N209A/I212A/N215A with mCGI-58, we observed a similar Lys/FT/W/EI/EII pattern compared to mATGL288/mCGI-58 WT suggesting that all mutant proteins bind to mATGL288 with similar efficiency.

**Figure 6.**
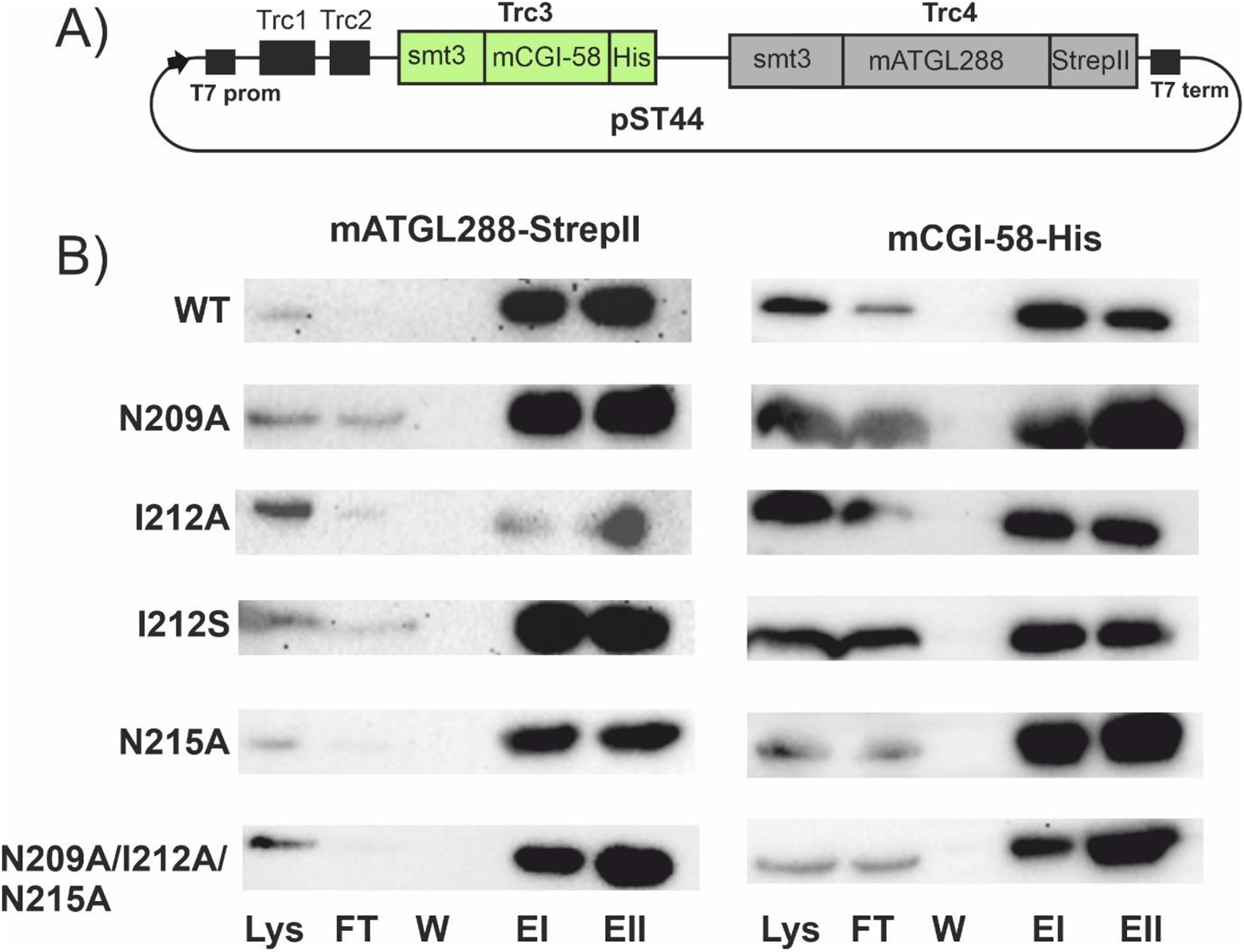
Co-purification of mATGL288 variants with mCGI-58. mATGL288 variants N209A, I212A, I212S, N215A and N209A/I212A/N215A with ablated co-activation are still able to interact with mCGI-58. A) Schematic overview of the polycistronic pST44 vector system. Cassette Trc3 codes for SMT3-mCGI-58 variants with C-terminal His-tags, whereas cassette Trc4 codes for SMT3-mATGL288 with a C-terminal Strep-II tag. B) Immunoblot analysis of the affinity purification experiments from the co-expression of the pST44 vector coding different mATGL288 mutants and mCGI-58. mATGL288 variants were detected with anti-StrepII antibody (SMT3-mATGL288, left) whereas mCGI-58 was detected with the anti-His antibody (right). The presence of proteins was monitored in fractions loaded onto the column (Lys), flow through (FT), last column wash (W) and two elution (EI and EII) fractions.

### The *in silico* generated 3D model of the ATGL/CGI-58 complex supports the general interaction involving the region spanned by N209 – N215 of ATGL and R228, G328, D334 of CGI-58

Using AlphaFold, we generated an *in silico* model of the ATGL/CGI-58 complex (Figure 7). For our analysis, we mapped the positions of N209, I212 and N215 of ATGL identified in this study and residues R228, G328, D334 of CGI-58 identified in previous work [16,35]. As seen in Figure 7A-7D, the interface between ATGL (truncated at residue 260 in this panel) and full-length mCGI-58 spans over large, yet continuous surface area of both proteins. I212 is central in a β-strand of a short two-stranded antiparallel β-sheet (E204-V207, T210-N213) at the far-end of the catalytic site in the predicted substrate binding pocket. N209 and N215 are located at the hinges of this β-strand. Substitution to of potential bidentate residues asparagine to alanine might change the flexibility of these hinges or be directly involved in changing protein-protein interaction (Figures 3, 4, 7). Importantly, data from experimental mutagenesis studies agreed very well with the calculated protein-protein interfaces. At the current stage of modelling, detailed analysis (e.g., with respect to inter-molecular H-bonding, formation of salt bridges) or fine-tuned modelling of the protein-protein interaction interface requires additional data, preferably from an experimentally determined structure (Figure 7B). The analysis of the C-terminal region of ATGL is challenging, since large parts of the C-terminal half are modelled with very low, low and at best medium confidences scores. Accordingly, the C-terminal half might adopt different secondary structures or different spatial positions in the physiological environment (e.g., the full LDs decorated with different LD associated proteins, Figure 7C). Next, we generated 3D models of truncated WT ATGL and the N209A/I212A/A215A ATGL variant. The triple variant exhibits large conformational rearrangements of region D197-L216-comprising a long loop and a short β-sheet in WT ATGL – by forming a loop and a long α-helix (Figure 7E). This region coincides with the region of high conservational variability (Figure 1) and low to medium confidence scores for structure prediction. It is interesting to note, that previous 3D homology modeling of the WT sequence of ATGL had an α-helix predicted for N209-L226, while the PNPLA domain (resides I10-K179) was essentially identical [41,42]. When we modeled the N209A/I212A/A215A ATGL/CGI-58 complex, the overall complex looked similar, however with some small changes in the interface. The model confidence for AlphaFold-Multimer models is described as an intrinsic score pTM and interface score ipTM [43]. The truncated WT ATGL/CGI-58 model exhibited a combined pTM + ipTM of 0.78, while the triple variant truncated ATGL/CGI-58 model, which experienced significant conformational changes in the loop region at the interface, had a score of 0.68. These results suggest a weaker interaction between the two proteins. Nevertheless, the score remains relatively high, indicating a reasonably strong interaction. There is an overall breathing motion and shifting of the co-activator CGI-58 to accommodate the longer α-helix and the lack of the short β-sheet of the ATGL variant (Figure 7F). Together, these predictions indicate conformational flexibility beyond the PNPLA-domain and awaits further experimental insights of ATGL by itself and in complex with the co-activator CGI-58.

**Figure 7.**
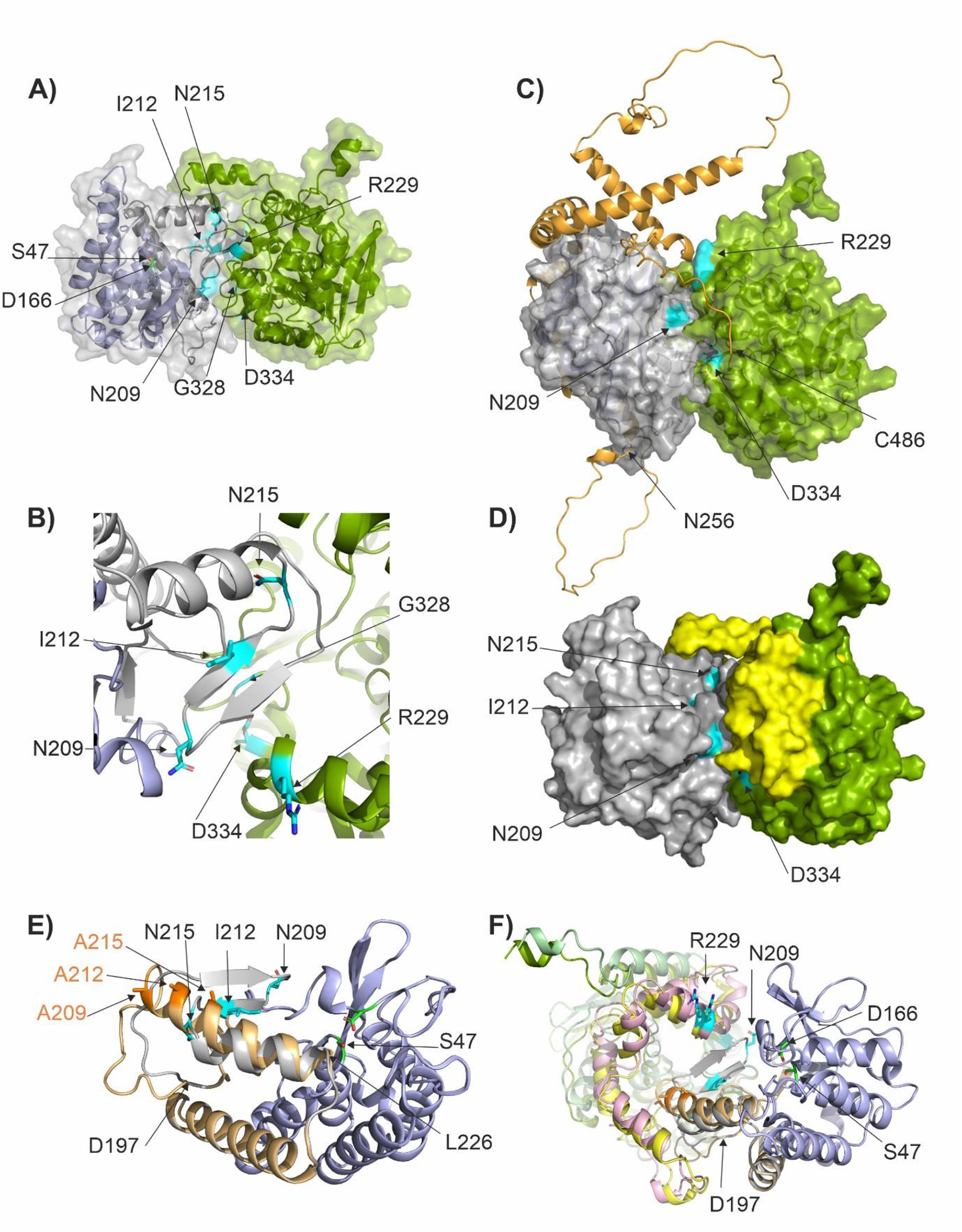
AlphaFold model of the ATGL/CGI-58 protein-protein complex. A-D) 3D models of mouse ATGL and mouse CGI-58 depicting important residues for activity and co-activation. The 3D structure of mATGL (M1-N259 in (A), M1-C486 in (C)) is in cartoon representation, the annotated PNPLA-domain (I10-L178), which is predominantly predicted with very high confidence, is colored light blue; residues M1-N9 and K179-L254 are in gray. Active site residues (S47, D166) are depicted as green sticks. mCGI-58 (G18-D351) is in pea-green cartoon representation. The transparent surface of both proteins is depicted in gray and pea-green, to highlight the extensive predicted protein-protein interaction surface. N209, I212 and N215 of ATGL as well as amino acids R299, G328 and D334 of CGI-58 are highlighted in cyan. B) Close-up view of the predicted interaction surface with N209, I212 and N215 of ATGL and amino acids R299, G328 and D334 of CGI-58 in cyan stick representation. C) 3D model of the ATGL/CGI-58 complex including the C-terminal half (N256-C486, colored in orange) of ATGL that is difficult to model with high confidence. D) Surface representation of the 3D model of the truncated ATGL/CGI-58 complex. The αβ-hydrolase core of CGI-58 is colored pea-green as in A-C, yet the cap region of CGI-58 (P180-M279) is colored in yellow. E) Overlay of the 3D models of truncated WT ATGL with the N209A/I212A/N215A ATGL variant. Regions predominantly predicted with very high confidence, are colored light blue; whereas regions K179-L254 with lower confidence in the prediction are in gray and sand for WT ATGL and N209A/I212A/N215A ATGL, respectively. ATGL is in similar orientation as in Figures 1B, 1C and 7F. F) Overlay of the complexes of WT ATGL/CGI-58 and that of the variant N209A/I212A/A215A ATGL with CGI-58. WT ATGL is colored light blue and gray; N209A/I212A/A215A ATGL is also in light blue, yet residues K179-L254 are in sand; the side chains A209, I212A and A 215 are depicted as orange sticks. When CGI-58 is in complex with WT ATGL, it is colored as in (B), whereas CGI-58 in complex with N209A/I212A/N215A ATGL is colored in light green for the core and pink for the cap, respectively.

## Discussion

Intracellular lipolysis is a crucial metabolic process in energy homeostasis and diligent balance of its regulation leads to metabolic equilibrium. CGI-58 is a crucial regulator of ATGL activity, but the mechanism by which CGI-58 co-activates ATGL remains elusive. It is not known if binding of mCGI-58 affects the conformation of ATGL, facilitates substrate presentation, or increases the lipolytic activity of ATGL by removing reaction products from the active site – questions similar to the unknowns discussed for the G0S2-mediated inhibition mechanism of ATGL [31]. We employed mutagenesis studies to identify specific amino acids in ATGL which are required for its co-activation by CGI-58. The sites for amino acid exchanges were selected based on evolutionary conservation and surface exposure. For most experiments, we used a shortened version of mouse ATGL, mATGL288, in combination with mouse full-length CGI-58 (mCGI-58) expressed in a bacterial expression system. Doing so, we could identify amino acids exchanges that resulted in complete loss of ATGL activity and exchanges leading to similar basal activity and activability by CGI-58 as observed for WT ATGL. Importantly, we identified the region N209-N215 of mouse ATGL to play an essential role in mCGI-58-mediated co-activation of ATGL (Figure 3, Figure 4). N209, I212 and N215 were recognized as residues with low evolutionary conservation (Figure 1) and they turned out to be crucial residues for co-activation based on *in vitro* assays (Figure 3). AlphaFold modeling revealed that these residues were also central for the mATGL288/mCGI-58 interaction in the interface of protein contacts (Figure 7). The findings suggest that the N209-N215 region is not directly involved in executing the hydrolysis reaction, but mediates CGI-58 co-activation of ATGL.

The comparison of ATGL variants from bacterial and mammalian expression system demonstrates that post-translational modifications are not required for the co-activation of ATGL by mCGI-58 (Figure 5). Furthermore, the comparison of full-length versus C-terminally truncated variants indicate that the full-length variants are slightly more tolerant towards single amino-acid exchanges. This is very similar to our previous observations on full length and truncated ATGL inhibition by variants of G0S2 [36]. The 254 N-terminal amino acids of mouse ATGL comprise the minimal domain that can be activated by mCGI-58 and inhibited by G0S2 [10]. Supposedly, additional regions within the C-terminal half of ATGL together with other factors either affect the protein-protein interaction, or provoke conformational changes in the lipase, the co-activator or the LD-associated TG substrates. ATGL variants associated with neutral lipid storage disease (NLSDM) are either enzymatically inactive proteins localizing to LDs or active TG hydrolases lacking LD localization [44]. Deletion of 220 amino acids from the C-terminus of human ATGL increases its interaction and activation by CGI-58 *in vitro*, in spite of defective LD localization *in vivo* in cultured cells [44]. This finding indicates that the C-terminal region of ATGL is required for its targeting to LDs and plays an important (auto)regulatory role that requires further characterization.

When we modelled the structure of the ATGL/CGI-58 complex, we found that large surfaces of both proteins, ATGL and CGI-58, are predicted to be involved in protein-protein contacts. Plotting N209, I212 and N215 on our 3D model of ATGL places these residues on an extension from the highly conserved PNPLA (I10-K179) domain on one face of the protein. The predicted interface of ATGL is completely in line with the herein presented and previous experimental results. Furthermore, the predicted interface region of CGI-58 matches well with previously published data: The Trp-rich N-terminal region is not predicted to be involved in the interface, which is in agreement with its annotated role in serving as LD localization anchor [33]. Both, the α/β-hydrolase core domain and the cap region (P180-M279, colored yellow in Figure 7D, 7F) of CGI-58 are predicted to be involved in formation of a large protein-protein interaction surface. The extent of the surface might also explain, why the single and even triple amino acid exchanges did not totally abolish the interaction in co-expression and co-purification assays (Figure 6). In the modeled ATGL/CGI-58 complex, the far end of the substrate binding cavity involving N209-N215 seamlessly merges into the pocket between the cap and the core of CGI-58. Previous studies had also demonstrated that CGI-58 interacts with fatty acid binding protein 4 (FABP4), which further promotes ATGL-mediated lipolysis and indicates the existence of an ATGL/CGI-58/FABP4 complex [45]. In the model of the binary ATGL/CGI-58 complex, a direct transfer of hydrophobic ligands (substrates or products) from ATGL to CGI-58 appears possible (Figure 7). AlphaFold-modeling of the binary CGI-58/FABP4 complex predicts that the helical portal region of FABP4 interacts with the cap region of CGI-58. This region of FABP4 has been shown experimentally to be involved in the FABP4/CGI-58 interaction using biochemical and biophysical methods [45].

In the absence of experimental 3D structures for ATGL and ATGL/CGI-58 complexes, our results represent a significant advance in our understanding of the protein-protein complex formed by ATGL/CGI-58 and provides novel insights into ATGL co-activation.

## Materials and Methods

### Materials

If not stated otherwise, chemicals were obtained from Merck (Darmstadt, Germany) or Carl Roth GmbH (Karlsruhe, Germany); columns for protein purification were obtained from Cytiva (formerly GE Healthcare Life Sciences (Uppsala, Sweden)). [9,10-^3^H(N)]-triolein was obtained from PerkinElmer Life Sciences (Waltham, MA, USA). Pierce™ Unstained Protein MW Marker from Thermo Scientific™ was used as size marker for SDS-PAGE gels. Blue Prestained protein standard broad range marker from New England BioLabs was used for SDS-PAGE and for Immuno Blotting. Disruption of cells was carried out using a homogenizer (SONOPLUS ultrasonic homogenizer HD 2070, Bandelin, Berlin, Germany).

### Site-directed mutagenesis of bacterial expression vectors

The WT variants coding for *Mus musculus* ATGL (UniProt Accession: Q8BJ56) in two different expression vectors namely pST44 and pcDNA4/HisMaxC (Thermo Fisher Scientific, Waltham, USA) were described in [46]. The primers listed in Table 1 and 2 were designed to introduce point mutations in the WT variants using the Q5® site-directed mutagenesis kit (New England BioLabs, Ipswich, USA). All variants were verified by Sanger sequencing (Microsynth, Balgach, Switzerland).

**Table 1:**
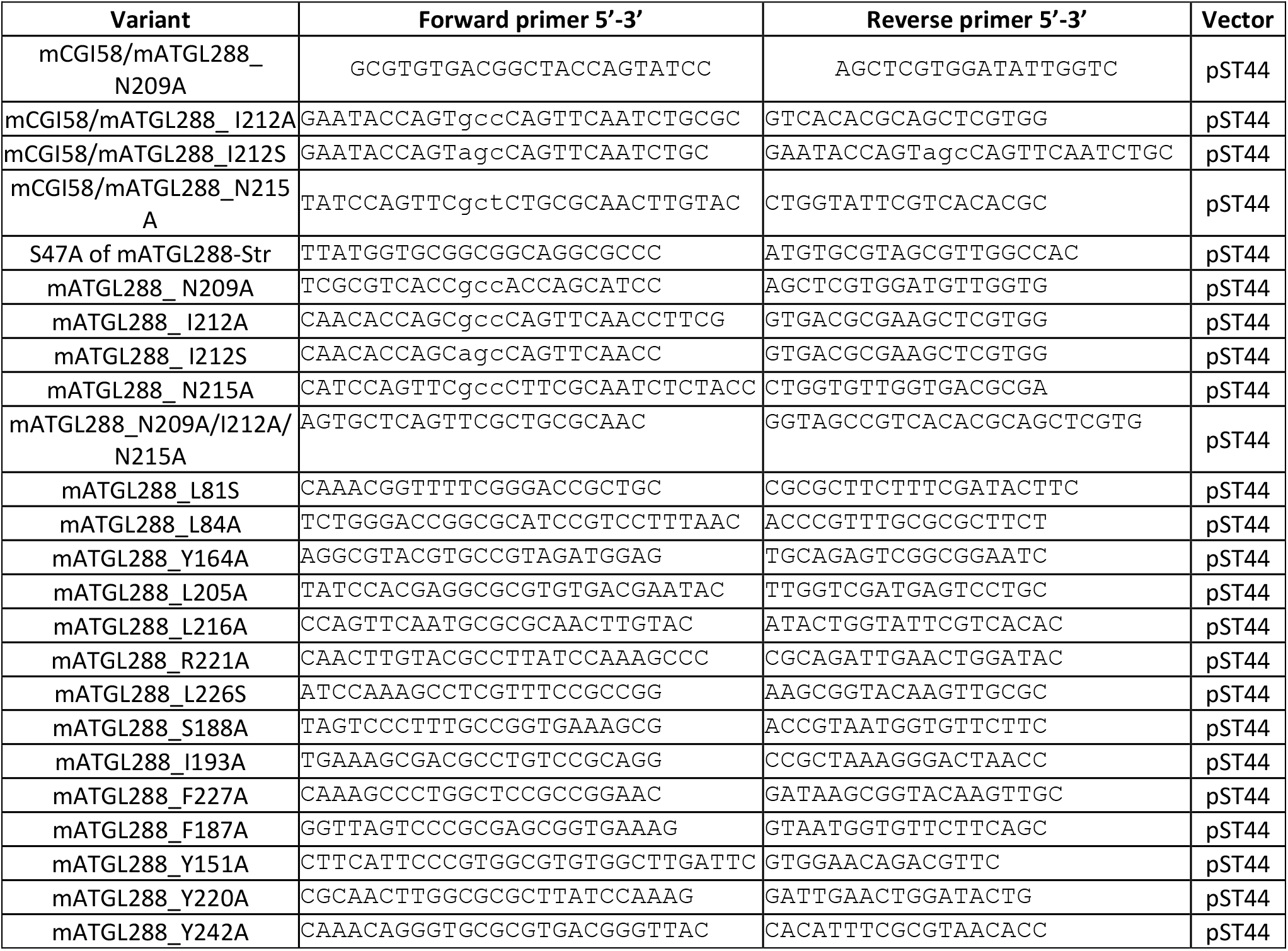
Primers for point mutations in mATGL288 and for mCGI-58/mATGL288.

### Site-directed mutagenesis of eukaryotic expression vectors

**Table 2:**
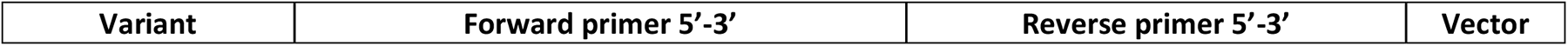
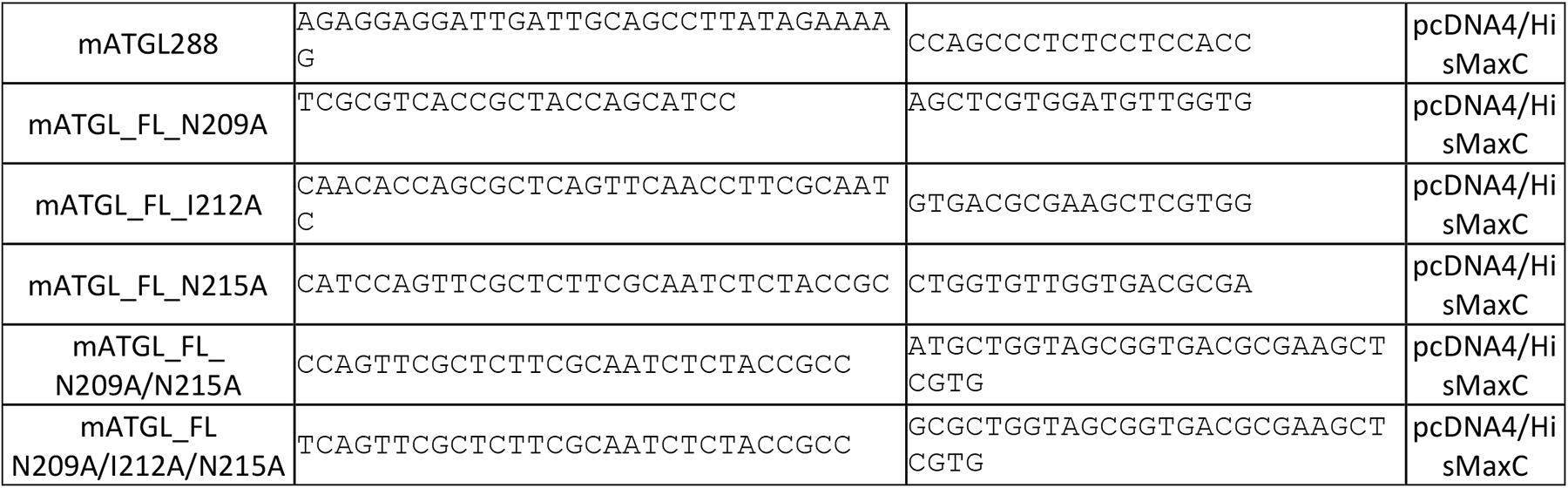
Primers for mATGL truncation and point mutations in eukaryotic expression vector pcDNA4/HisMaxC.

### Cloning of pST44-Trc3-mCGI58-His/Trc4-mATGL288-Str

The binary pST44-Trc3-mCGI58-His/Trc4-mATGL-Str construct was cloned by linearization of pST44-Trc3-smt3-TEV-GGG-mCGI58-His with BspEI and MluI as well as linearization of pST44-Trc4-smt3-TEV-GGG-mATGL288-Str with SacI and KpnI. The inserts required for ligation into translational cassette three and four of the linearized pST44 vectors, were generated by complementary cutting of pST44-Trc3-smt3-TEV-GGG-mCGI58-His with SacI and KpnI and cutting of pST44-Trc4-smt3-TEV-GGG-mATGL288-Str with BspEI and MluI. Ligation reaction of the binary polycistronic pST44-Trc3-mCGI58-His/Trc4-mATGL-Str was performed with T4 DNA Ligase.

### Bacterial expression of recombinant mCGI-58, ATGL-proteins, mATGL/mCGI-58 complexes and preparation of bacterial cell extracts

Expression of mATGL288 and mCGI-58 have been described before [5]. Expression of mATGL288 protein and single amino acids exchange variants were performed in *E. coli* ArcticExpress (DE3) cells (Agilent Technologies, Santa Clara, CA, USA) similar as previously described [5]. Expression of the complexes mATGL288/mCGI-58, mATGL_N209A/mCGI-58, mATGL_I212A/mCGI-58, mATGL_N215A/mCGI-58 and mATGL_N209A/I212A/N215A/mCGI-58 complexes were also performed in *E. coli* ArcticExpress (DE3) cells. A day culture (LB media supplemented with 100 μg/ml ampicillin, 20 μg/ml gentamycin and 2% glucose) was started and incubated for 8 h at 37 °C. The day culture was transferred (1:50) to a preculture (with the same supplements as the day culture) for 8 h growth at 37 °C. The main culture (LB media +0.5% glucose) was inoculated (1:10) with preculture and grown at 30 °C until it reached an OD_600_ of 0.5. Cells were then induced with 0.1 mM of isopropyl-β-d-thiogalactopyranoside (IPTG) and harvested after 24 h of expression at 10 °C. After harvesting the cells were frozen and stored at -20 °C before further usage. All mATGL288 variants were treated equally. Prior the activity assays, the cells for mATGL288 variants were resuspended in sucrose solution (250 mM sucrose, 1 mM ethylenediaminetetraacetic acid (EDTA), 1 mM 1,4-dithiothreitol (DTT), 20 μg/ml leupeptine, 2 μg/ml antipain, 1 μg/ml pepstatin, pH 7.0) and disrupted by sonication on ice. After centrifugation at 15000 × *g* for 20 min at 4 °C supernatants were collected. Total protein concentrations of the cell extracts were determined using the Bio-Rad Protein Assay Kit according to the manufacturer’s instructions (Bio-Rad Laboratories, Hercules, CA, USA) using BSA as standard.

### Co-purification of mATGL/mCGI-58 complexes

mATGL288 was co-expressed with mCGI-58 in ArcticExpress (DE3) cells. A 1 l cell pellet was thawed on ice and resuspended in 20 ml Lysis Buffer (100 mM K_2_HPO_4_/KH_2_PO_4_ pH 7.5, 100 mM KCl, 30 mM imidazole, 10% glycerol, 0.1% IGEPAL CA-630, 1 mM TCEP, 10 mM ATP, 10 mM MgCl_2_) supplemented with 20 μl of 1000X protease inhibitor (PI) stock solution (1.5 mM pepstatin A, 3.3 mM antipain and 43 mM leupeptin). After cell lysis via sonication (10 minutes, cycle 5, 50% power) and centrifugation (40 minutes, 20,000 x g, 4°C), the cell lysate was filtered using a 0.45 μm syringe filter. The chromatography steps were carried out using the ÄKTA Avant 25. As a first step, the lysate was loaded onto a 1 ml StrepTrap HP column (GE Healthcare Life Sciences, Buckinghamshire, UK) that was equilibrated with Lysis buffer.

The column was washed with 15 CV of Lysis Buffer, followed by 20 CV of Nickel-Wash Buffer (100 mM K_2_HPO_4_/KH_2_PO_4_, pH 7.5, 500 mM KCl, 30 mM imidazole, 10% glycerol, 1 mM TCEP, 10 mM ATP, 10 mM MgCl_2_) and 20 CV of Strep-Wash Buffer (100 mM K_2_HPO_4_/KH_2_PO_4_, pH 7.5, 100 mM KCl, 10% glycerol, 1 mM TCEP). The proteins were eluted using 10 CV of a 100% Strep-Elution Buffer (100 mM K_2_HPO_4_/KH_2_PO_4_, pH 7.5, 100 mM KCl, 10% glycerol, 1 mM TCEP and 10 mM desthiobiotin). Elution fractions were subjected to 12% SDS-PAGE to verify the presence of proteins. Samples with highest amounts of proteins were used for immunoblotting, using Strep-antibody against mATGL and His-antibody against mCGI-58. Of each variant a load (L), wash (W) and elution (E) sample were subjected to SDS-PAGE. “Load” stands for the cleared cell lysate which was loaded onto the column and “wash” is the last wash fraction taken before the elution.

### Protein expression in Expi293F^TM^ cells and preparation of cell-extracts for TGH assays

His-mATGL and its point mutants were recombinantly expressed in Expi293F^TM^ cells (Thermo Fisher Scientific, Waltham, USA). The cells were cultivated in Expi Expression Medium (Thermo Fisher Scientific, Waltham, USA) at standard conditions (37°C, 95% humidified atmosphere, 7% CO_2_). Cell density was determined in a CASY Cell Counter and Analyzer System (OMNI Life Science, Bremen, Germany). Cells in a total culture volume of 10 ml were transfected with 10 µg of plasmid DNA using the ExpiFectamineTM 293 Transfection Kit (Thermo Fisher Scientific) according to the manufacturer’s instructions. Two days after transfection, cells were harvested by centrifugation at 500 x g and 4°C for 5 min. The cell pellet was washed twice with PBS (137 mM NaCl, 2.7 mM KCl, 10 mM Na_2_HPO_4_, 1.8 mM KH_2_PO_4_). Cells were disrupted immediately after harvesting by sonication (Sonopuls GM3100 equipped with a MS-72 tip; Bandelin, Berlin, Germany) in sucrose solution (250 mM sucrose, 1 mM EDTA, 1 mM DTT, 20 μg/ml leupeptin, 2 μg/ml antipain and 1 μg/ml pepstatin) on ice. The homogenate was centrifuged at 1,000 x g and 4°C for 20 min. The post-nuclear fraction was collected and protein concentration was determined by the Bradford protein assay (Bio-Rad Laboratories, Hercules, USA) using bovine serum albumin (BSA) as standard.

### Purification of recombinant mouse mCGI-58

Purification of His_6_-smt3-mCGI-58 was performed via immobilized metal ion affinity chromatography as previously described [5]. The concentration of the purified protein was determined via absorption at 280 nm.

### TG Hydrolase Assay

TG hydrolase activity assays was performed with some minor modifications as described elsewhere [5,10,36]. To screen for the stimulation capacity of various mATGL288 variants, 25 μg protein of cell lysates containing ATGL were incubated together and 1 μg purified mCGI-58 in a total volume of 25 μl sucrose solution (250mM sucrose, 1mM EDTA, 1mM DTT, pH 7.0) with 25 μl ^3^H-triolein substrate. Empty uninduced ArcticExpress (DE3) cells were used as a negative control. TG substrate was prepared with 1.67 mM triolein, 25 μCi/ml [9,10-^3^H(N)]-triolein (PerkinElmer Life Sciences, Waltham, MA, US) and 188 μM L-α-phosphatidylcholine: L-α-phosphatidylinositol PC:PI (3:1) (Merck, Darmstadt, DE). 1.5 ml of 0.1 M potassium phosphate buffer, pH 7.0, was added and the mixture was subsequently sonicated. After sonication, fatty acid (FA) free BSA (Merck, Darmstadt, DE) was added to a final concentration of 5%. As a blank measurement, 25 μl of sucrose solution were mixed with 25 μl of substrate solution (50 μl of reaction mixture). The protein-substrate mix was then incubated for 1h at 37°C shaking. The reaction was terminated by adding 650 μl of methanol:chloroform:heptane (10:9:7) and 200 μl of 100 mM potassium carbonate buffer pH 10.5 (adjusted with boric acid). The radioactivity in 100 μl of the upper phase was determined by liquid scintillation counting. FA extraction was performed by mixing the samples vigorously for 5 sec followed by centrifugation at 2,400 x *g* at room temperature for 10 min. An aliquot of 100 μl of the upper aqueous phase was transferred into scintillation vials containing 2 ml of scintillation cocktail (Roth). Radioactivity was determined by liquid scintillation counting using a MicroBeta Microplate Counter (PerkinElmer, Waltham, Massachusetts). Specific substrate activities were in the range of 1000-2000 cpm/nmol. Counts generated from blank measurements were subtracted and the rates of TG hydrolase activity, presented as nmol of released FA per hour and milligram of protein, were calculated according to Schweiger *et al*. [47]. Calculations of EC_50_ values were done using approximation with sigmoidal function in Microsoft Excel.

### Immunoblotting

Bacterial lysates/purified proteins containing ATGL variants with point mutations and samples of mATGL288/mCGI-58 after co-purification were loaded and run on a denaturing 12% SDS-PAGE gel and transferred to PVDF membrane (ROTH, pore size 0.45 μm) for 90 minutes at constant current of 220 mA. After the transfer, membranes were treated using standard protocols for Anti-His or Anti-Strep immunoblotting with minor modifications. For both anti-His and anti-Strep immunoblotting, standard Tris buffered saline buffer (TBS, 50 mM Tris-HCl pH 7.5, 150 mM NaCl) and tris buffered saline tween buffer (TBST, 50mM Tris-HCl pH 7.5, 150mM NaCl, 0.2% v/v Tween 20) were used. For anti-His immunoblotting all membranes (mATGL288_WT/mCGI-58, mATGL288_N209A/mCGI-58, mATGL288_I212A/mCGI-58, mATGL288_I212S/mCGI-58, mATGL288_N215A/mCGI-58, mATGL288_N209A/I212A/N215A/mCGI-58) were blocked with 3% BSA in TBS buffer for 90 minutes at room temperature. Afterwards they were washed 2×10 minutes in TBST and 1×10 minutes in TBS buffers accordingly. The PentaHis antibody (Qiagen, Düsseldorf, DE) against His-Tagged proteins was diluted 1:5000 in TBS buffer containing 3% BSA and the membrane was incubated overnight at 4°C followed by 2×10 min washing steps in TBST and 1×10 min in TBS buffers. The secondary antibody ECL anti-mouse IgG (GE Healthcare Life Sciences, Buckinghamshire, UK), diluted 1:5000 in TBS buffer containing 5% milk powder was applied to the membrane for 90 minutes at room-temperature. For anti-Strep immunoblotting all membranes (mATGL288_WT/mCGI-58, mATGL288_N209A/mCGI-58, mATGL288_I212A/mCGI-58, mATGL288_I212S/mCGI-58, mATGL288_N215A/mCGI-58,mATGL288_N209A/I212A/N215A/mCGI-58) were blocked with 3% BSA in TBST buffer containing 10% milk powder for 90 minutes at room temperature, followed by 2×10 minutes washing steps in TBST buffer. StrepMAB-Classic mouse antibody (IBA Lifesciences, Goettingen, DE) was diluted 1:2000 in TBST buffer containing 5% milk powder and the membrane was incubated overnight at 4 °C. Next morning the membrane was washed 2×10 min in TBST buffer. The secondary antibody ECL anti-mouse IgG (GE Healthcare Life Sciences, Buckinghamshire, UK), diluted 1:5000 in TBST containing 5% milk powder was applied to the membrane for 90 minutes at room temperature. Both anti-His and anti-Strep immunoblots were developed using Amersham ECL immunoblotting reagents (GE Healthcare Life Sciences, Buckinghamshire, UK).

### Modelling of ATGL and the ATGL/CGI-58 complex using AlphaFold

ATGL and hetero multimers of ATGL with CGI-58 (ABHD5) were calculated using AlphaFold multimer. The predictions were based on either full-length protein from mouse (ATGL M1-C486; CGI-58 M1-351) or a truncated form of ATGL with a C-terminal truncation (ATGL M1-D264). Despite the differences, both predictions resulted in the identification of the same interaction interface. Furthermore, the same result was obtained when both combinations were predicted using AlphaFold monomer mode with an unstructured 50x glycine linker between the two different proteins. Predictions were either calculated with AlphaFold v2.1.2 full installation on a local workstation with following specifications: Nvidia GeForce RTX 3090 24 GB, AMD Ryzen Thread Ripper 3975WX using 48 cores and 192 GB RAM or with AlphaFold v2.3.1 and local ColabFold v1.5.2 [48] installation using a workstation with Nvidia GeForce RTX 3090 24 GB, AMD Ryzen 9 5900X using 12 cores and 64 GB RAM.

## Abbreviations

ATGL: adipose triglyceride lipase
HILPDA: Hypoxia inducible gene-2
G0S2: G0/G1 switch gene 2
LD: lipid droplet
CGI-58: comparative gene identification-58
FA: fatty acid
IPTG: isopropyl β-D-1-thiogalactopyranoside
PNPLA: patatin-like phospholipase
SDS: sodium dodecyl sulfate, TG, triacylglycerol
WT: wild-type.

## Acknowledgements

This study was supported by a grant by the Austrian Fonds zur Förderung der Wissenschaftlichen Forschung (SFB F73 Lipid Hydrolysis to RZ and MO) and the doc.fund projects Molecular Metabolism (DOC 50 to MO) and Biomolecular Structure and Interactions (DOC 130 to MO and TPL) funded by the Austrian Science Fund FWF, Land Steiermark, the City of Graz.

